# Long-range neural pathways for octopus chemotactile processing revealed from periphery-to-brain by centimeter-field microCT

**DOI:** 10.64898/2026.01.02.697341

**Authors:** Andrew Sugarman, Daniel Vanselow, David Northover, Stephen L. Senft, Carolyn Zaino, Maksim A. Yakovlev, Jessica Christ, Justin D. Silverman, Mee S. Ngu, Khai C. Ang, Steve Wang, Wen-Sung Chung, Patrick La Riviere, Roger T. Hanlon, Keith C. Cheng

**Affiliations:** Department of Pathology, Penn State College of Medicine, Hershey, Pennsylvania, USA, The Jake Gittlen Laboratories for Cancer Research, Penn State College of Medicine, Hershey, Pennsylvania, USA; Program in Bioinformatics and Genomics, Pennsylvania State University, State College, Pennsylvania, USA; Medical Scientist Training Program, Penn State College of Medicine, Hershey, Pennsylvania, USA; The Marine Biological Laboratory, Woods Hole, MA, 02543, USA; College of Information Sciences and Technology, Pennsylvania State University, State College, Pennsylvania, USA; Department of Statistics, Pennsylvania State University, State College, Pennsylvania, USA; Department of Medicine, Penn State College of Medicine, Hershey, Pennsylvania, USA; Institute for Computational and Data Science, Pennsylvania State University, State College, Pennsylvania, USA; Mobile Imaging Innovations, Inc., Palatine, Illinois, USA; School of the Environment, The University of Queensland, St Lucia, QLD, Australia; Department of Radiology, The Univ. of Chicago, USA; Molecular & Precision Medicine Program, Penn State College of Medicine, Hershey, Pennsylvania, USA

**Keywords:** Whole-organism phenotyping, digital organismal biology, histotomography, wide-field microCT, virtual 3D histology, sensory biology, distributed nervous system, nervous system architecture, organ system architecture, chemosensory nervous system, sucker neurocircuitry, arm-to-arm U tracts, intermediate longitudinal tracts

## Abstract

Understanding how nervous systems mediate responses to sensation requires whole-body maps of periphery-to-brain connections. Octopuses exemplify this challenge with distributed control of eight arms and hundreds of suckers, yet their long-range microanatomical wiring remains elusive due to limitations in microscopy. We extend histotomography (Ding et al. 2019), a form of soft tissue microCT customized for volumetric characterization of cells and tissues, to centimeter range with a custom micro-CT imaging system (Ding et al., 2019). With its 10-mm field of view and 0.7-µm isotropic voxels we created a high-resolution digital intact small octopus. This multi-tissue 3D blueprint enabled us to (i) elucidate previously uncharacterized chemotactile pathways from the suckers to the brain, (ii) discern subdivisions of the nerve ring connecting neighboring arms, and (iii) segment over 300 structures across organ systems at histology-like resolution. We release the labeled interactive digital specimen to facilitate collaborative whole-organism phenotyping as a practical foundation for *digital organismal biology*.

**ONE-SENTENCE SUMMARY:** *Whole-body 3D histology reveals neural and organ architecture throughout a small octopus*.

## INTRODUCTION

3D histological phenotyping across tissues and whole organisms such as octopus using existing technology has several challenges (Cheng et al., 2011, 2016). For example, light-sheet fluorescence microscopy (LSFM) achieves rapid volumetric imaging of large samples including live zebrafish, human tumor biopsies, mouse brains, and whole *Drosophila* (Daetwyler & Fiolka, 2023; Glaser et al., 2022; Mcconnell & Amos, 2018; Tian et al., 2024). However, its reliance on molecule-specific stains for contrast and limited penetration of visible light restricts the study of all cell types in single volumes of multimillimeter sized samples (Bürgers et al., 2019; Daetwyler & Fiolka, 2023; Mai et al., 2024). Serial section and block-face electron microscopy (SSEM) provide striking nanometer-range resolution in fixed tissue, enabling characterization of synapses and axonal connections, but such studies typically have been limited in sample volume to less than a cubic millimeter, are anisotropic, require labor intensive and destructive manipulation of the specimen, limiting whole-organism context (Hildebrand et al., 2017; Neacsu & Crook, 2024). High-field magnetic resonance and diffusion MRI offer non-destructive 3D anatomy but at voxels sizes of tens of microns, restricting the assessment of cellular-scale structures (Chung et al., 2020, 2022; Johnson et al., 2023).

X-ray micro computed tomography (microCT) provides a non-destructive 3D alternative at a detailed mesoscale, with synchrotron and some bench-top sources yielding isotropic, sub-micrometer voxels (Ding et al., 2019; Frost et al., 2023; Katsamenis et al., 2019; Yakovlev et al., 2022; Zehbe et al., 2009). Conventional microCT typically couples the scintillator to standard microscope objectives that achieve micrometer-scale spatial resolution over fields of view of <2mm. This gap has made cellular resolution imaging at the scale of whole organisms and large tissue samples impractical. Our lab addressed this problem by developing custom detectors, such as the expanded 5mm-wide FOV characterized by Yakovlev et. al in 2022 (Yakovlev et al., 2022). Here, we introduce a 10mm wide detector for micro-CT that brings micrometer resolution to the whole-organism scale, which we showcase with a detailed 3D examination of an intact hatchling octopus and a portion of an adult arm.

The octopus exhibits complex behaviors based on a distributed and convoluted nervous system, which exemplifies the challenge of identifying long-range connections throughout an entire animal. Their extraordinary degree of environmental touch and chemotactile sensing involves hundreds of suckers within and across 8 arms (Bidel et al., 2022, 2023; Chang & Hale, 2023; Fouke & Rhodes, 2020; Hanlon & Messenger, 2018; Kennedy et al., 2020; Kuuspalu et al., 2022; Sepela et al., 2025; Wells, 1978a). The octopus’ sensory and motor systems traverse its entire body with a robust nervous system of roughly 500 million neurons, 2/3 of which are arranged in the arms (Carls-Diamante, 2022; “Chapter 3: The Nervous System of The Arms,” 1971; Young, 1971). Detailed characterization of the long-distance sensorimotor pathways linking suckers and the central brain remains fragmented because methods capable of resolving microconnectivity are confined to small fields of view.

The combined FOV and resolution of our 10mm detector system enabled the first 3D volumetric reconstructions of an intact hatchling and adult arm with isotropic detail and with contrast resembling that of histology. Our choice of *Octopus bimaculoides* was driven by the size and complexity of its hatchling stage, which, aside from the reproductive system, contains fully developed organ systems and exhibits the sophisticated behaviors of adults (Figure S1) (Forsythe & Hanlon, 1988; Hanlon & Messenger, 2018). We segmented microanatomy across olfactory, visual, vascular, respiratory, and digestive systems, and most focally, the chemosensory nervous system. Neural characterizations revealed new aspects of touch- and taste-driven coordination, specifically, intermediate longitudinal nerve tracts (iLTs) – likely corresponding to the ‘bilateral longitudinal tracts’ recently identified in isolated arms (Olson & Ragsdale *biorxiv*, 2025) – that we follow from suckers into brain regions involved in sensory integration, learning and memory, and subdivisions of the nerve ring that likely mediate inter-arm communication (Olson & Ragsdale, 2025). We release the labeled 3D dataset as an open “digital specimen” to facilitate validation, discovery, and collaboration. This web-based resource (Maitin-Shepard, 2021) serves as a practical foundation for *digital organismal biology*, providing native architecture to ground future models linking structure to physiology, behavior and environment (Maitin-Shepard et al., 2021).

## RESULTS

### Whole-organism 3D histology via centimeter-field sub-micron CT

To facilitate multi-system, whole-organism understanding of microanatomy, we extended the field of view of microCT-based “3D histology”, *histotomography* (Ding, 2019) (Ding et al., 2019). The novel micro-CT detector system features a fixed-position scintillator, 12mm diameter lens FOV, 0.6 numerical aperture objective lens with a sCMOS camera that has a 10 x 7 mm viewing area and 0.7-micrometer pixels (Figure 1, Figure S2). The spatial resolution of this system as measured with a QRM Nanobar phantom was ∼1.6 micrometers in the projection domain and ∼2.3 micrometers in the reconstruction domain (Figure S3). The unprecedented combination of centimeter scale FOV and 0.7-micrometer isotropic voxel size of the reconstructed volume made practical the segmentation of every major organ system (Figure 1, Figure S4, Video S1 Tour), including the multiple tissues of the nervous system from skin to brain, without destructive sectioning or excessive reliance on artifact-inducing tomographic stitching (Figure 1, Figure S2) (Fransson et al., 2024; Schindelin et al., 2015; Schroeder et al., 2021; Vo et al., 2021).

**Figure 1:**
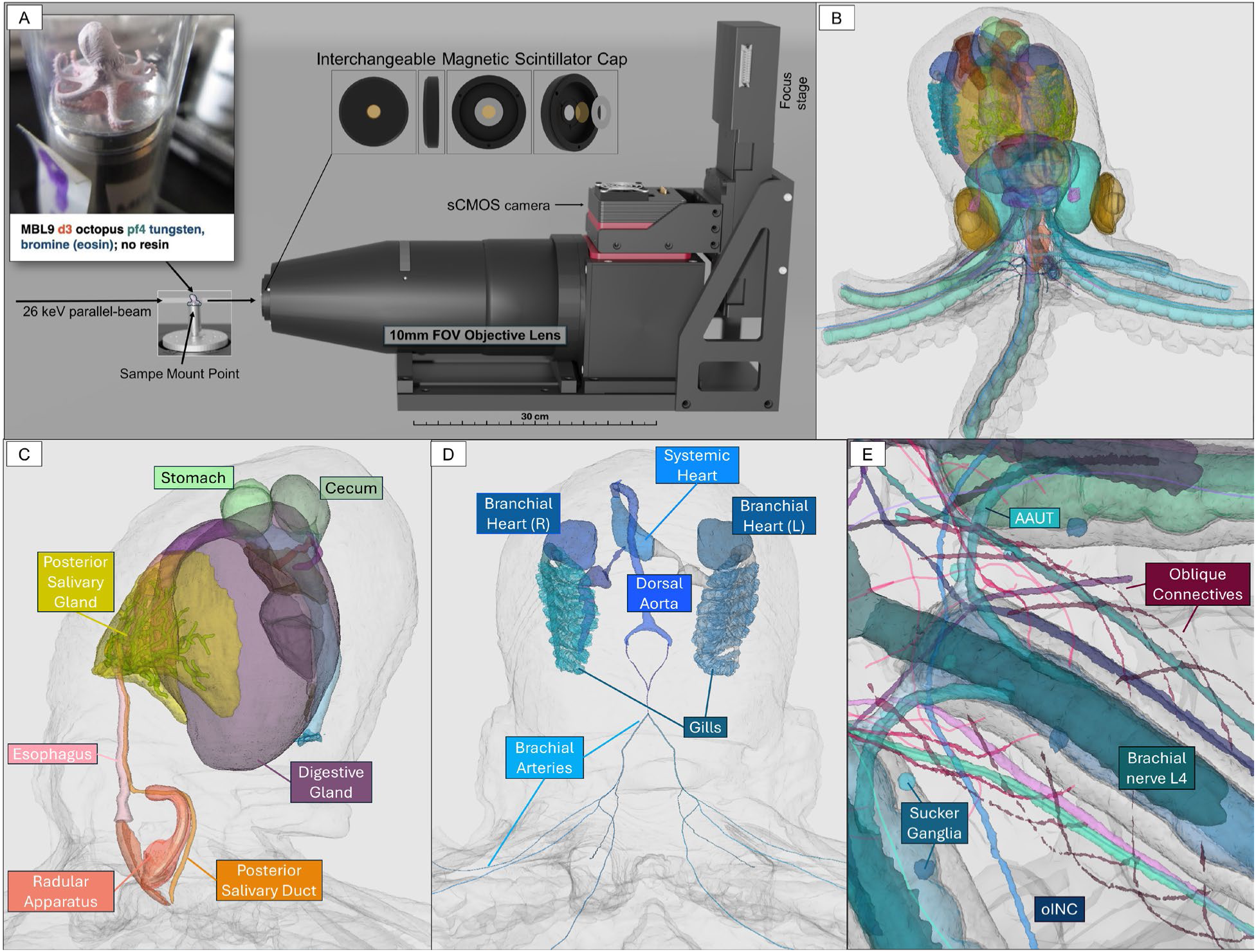
(A) Whole octopus histotomography enabled by custom wide-field detector. Centimeter-wide rotational projections of X-rays passing through the rotating formalin-fixed, metal-stained O. bimaculoides (pink) onto a scintillator (tan disc, (B)) were magnified by the custom lens and mirror onto a medium format sCMOS chip camera (B). (C) 3D Dragonfly of render the reconstructed hatchling octopus, with a spherical section removed virtually to reveal the long-range pathway of the brachial nerves extending from the arms in the periphery, through the nerve ring in the mantle, and into the central brain. 2D slices of tomographic reconstructions were used to segment and pseudocolor microanatomical structures (shown together, top left).Bottom panels (D) include the digestive, circulatory, whole nervous system, and its arm branches. (E) Snapshot taken from the peripheral nervous system, highlighting segmentations of the nerve ring (AAUT shown here), brachial nerve (BN), oblique connectives (OCs), and oral intramuscular nerve cords (oINCs).

To maximize image quality and resolution using this wide-field system, we utilized the high-flux and parallel beam geometry of Sector-2 Bending Magnet (2-BM) beamline X-rays at the Advanced Photon Source synchrotron of Argonne National Laboratory (Gürsoy et al., 2014; Nikitin et al., 2022). The intact *Octopus bimaculoides* hatchling was prepared in the Hanlon laboratory at the Marine Biological Laboratory with heavy-metal staining to increase image contrast (Ding et al., 2019; Katz et al., 2021; Metscher, 2009). To minimize perturbation of the native anatomy during sample preparation and signal attenuation during scanning, we did not embed the organism. Our detector system (10 mm wide, 7 mm tall) can accommodate the full height of our mounted specimen, but since the octopus exceeded the 2-BM beam height, two scans were used to capture the full height of the sample (Preibisch et al., 2009). The whole-organism image repository allowed us to pursue a function-driven interrogation of new aspects of the octopus’ widely distributed chemotactile pathway (Figures 2 and 3). The image includes the highest resolution 3D image of a whole nerve ring to date, within which subdivisions linking neighboring brachial (arm) nerves are apparent (Figure 4, Figure S5). The resolution achieved also allowed labeling of other prominent peripheral neural structures including series of oblique connectives and the arm-skipping oral intramuscular nerve cords (Figure 1E, Figure S6, S7, Video S2 INC 3D Video S3 OC 3D).

**Fig. 2:**
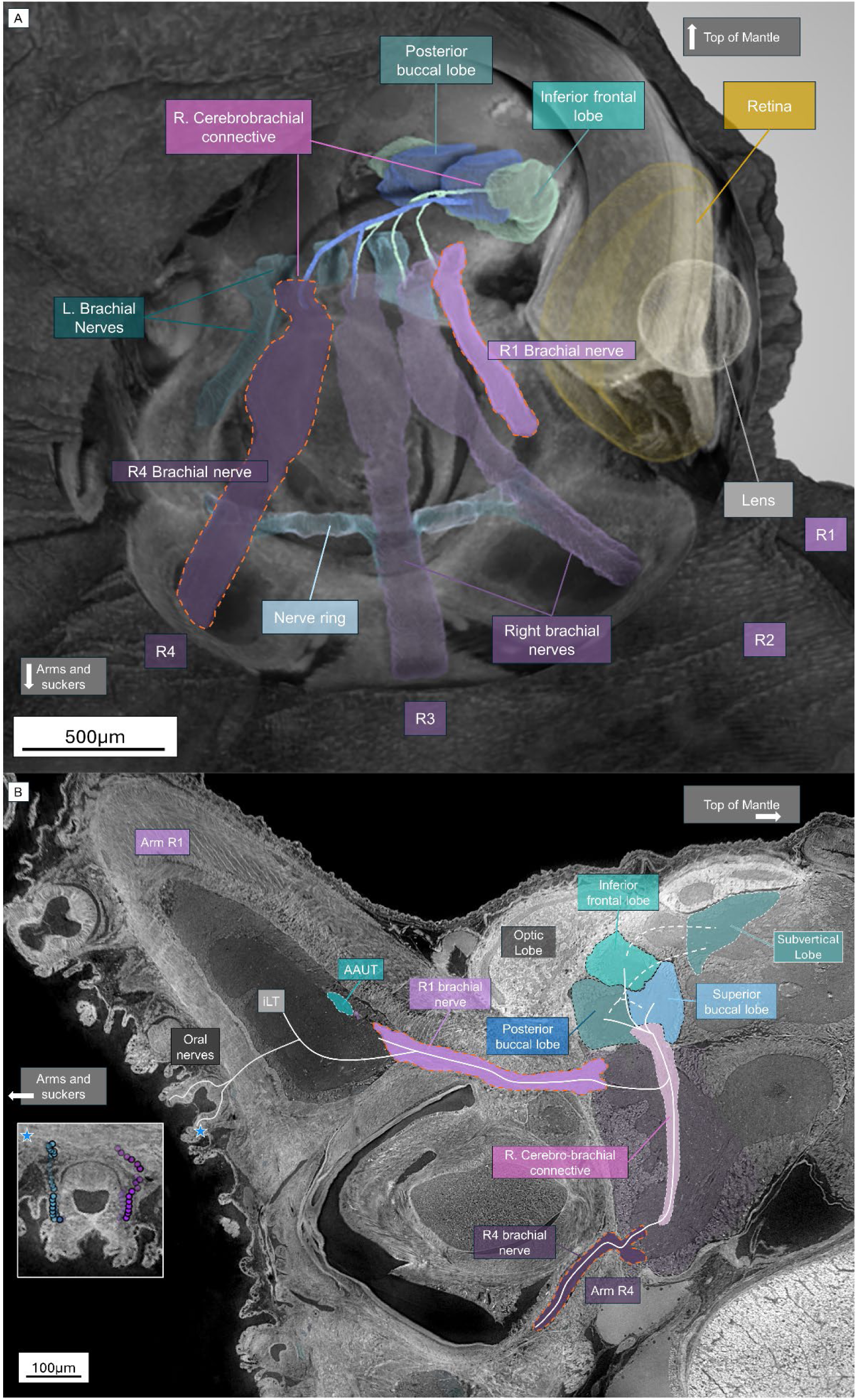
3D mapping of the long-range connections from sucker to brain in Octopus. (A) 3D render of hatchling reconstruction with spherical cutout illustrating the brachial nerves entering the body from the arms and projecting into the central brain. Each brachial nerve contributes to the portion of the cerebrobrachial connective that enters the inferior frontal lobe (shown here from the right 4 arms only for visual simplicity). Dotted line indicates contiguous brain regions (frontal lobes and subvertical lobe). (B) 2D sagittal cross section of the hatchling octopus from arm (left side) to brain (right side). Schematic lines illustrate continuous textures labeled connecting the oral nerves to the intermediate longitudinal tracts (see also Fig. 3) to the brachial nerves, which project to the cerebrobrachial connectives. The CBCs then project to the buccal lobe and the inferior frontal lobe (IFL). Dotted lines indicate connections between brain regions that exist but are not visible at the chosen 2D cross section.

**Fig. 3:**
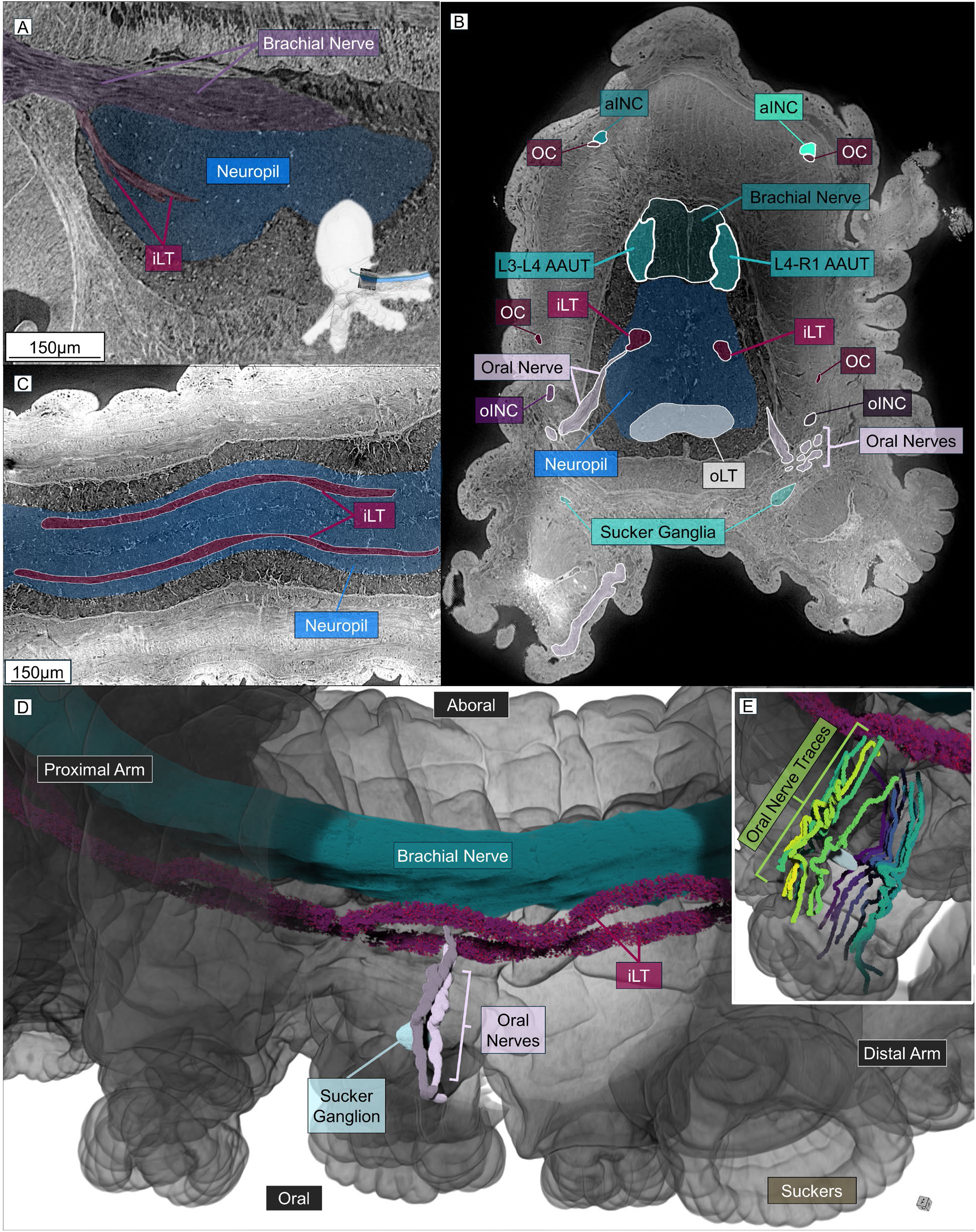
Intermediate longitudinal tracts (iLT) illustrate distinct pathway linking oral nerves from the suckers to the proximal brachial nerve. (A) Sagittal cross-section of proximal end of arm L4 within the hatchling octopus reconstruction, illustrating the texture we labeled as the iLT descending separately from the brachial nerve into the neuropil of the arm. (B) Transverse cross-section of the excised adult arm reconstruction illustrating the relative position of the iLTs within the neuropil and their intersection with the oral nerves. (C) Coronal cross-section of the excised adult arm reconstruction illustrating the paired iLT structures running parallel within the neuropil. (D) 3D rendering of the adult arm reconstruction with a cutout revealing the oral nerves, their intersection with the iLT, and the spatial positioning of the iLT and brachial nerve within the axial nerve cord (ANC). (E) Alternative angle illustrating the proximity and contact of oral nerves with the sucker ganglion. Segmented oral nerves are rendered in color, and additional oral nerve traces are rendered in situ.

**Figure 4:**
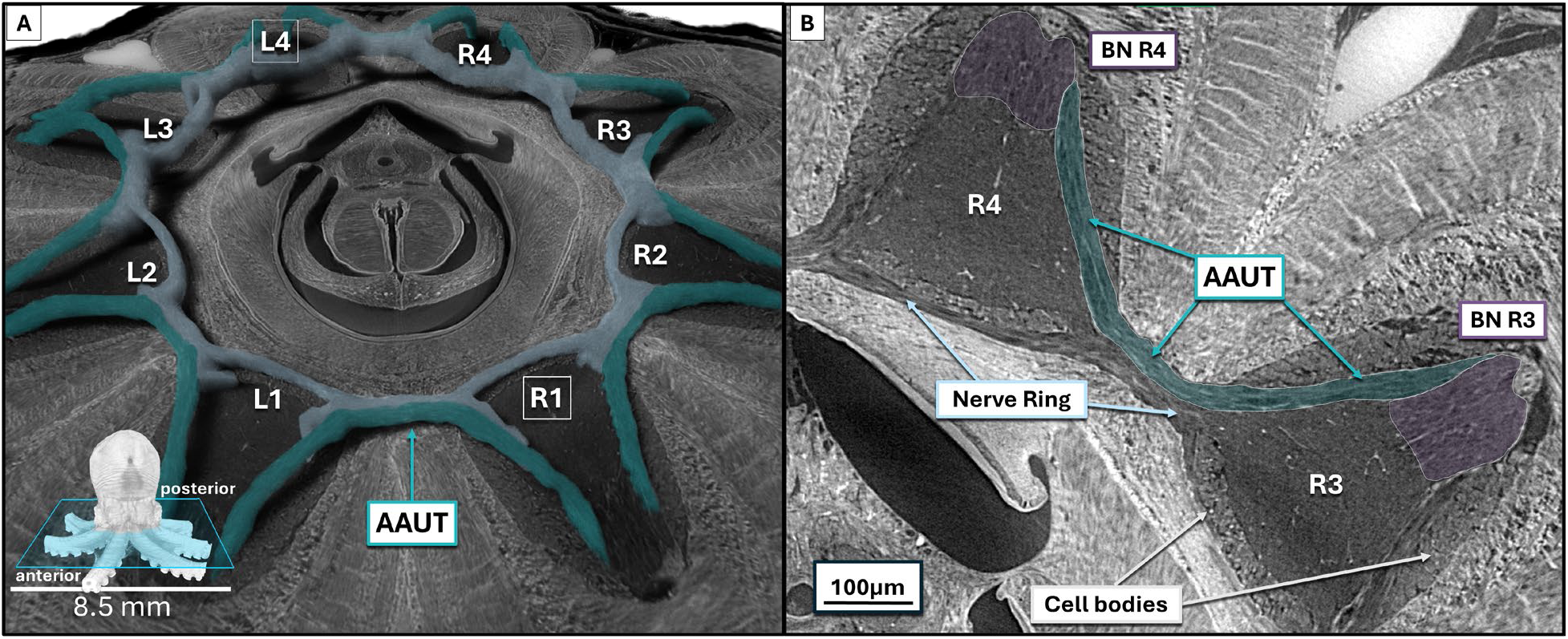
Subdivisions of the octopus nerve ring. A. Axial cross-section of the reconstructed octopus histotomogram at 0.7-micrometer resolution with nerve ring segmentation rendered in 3D. The green subdivision delineates what we refer to as the Arm-to-Arm U-Tract (AAUT), connecting the brachial nerves (BN) of neighboring arms via nerve fascicles that flow through each interbrachial commissure. The blue subdivision represents the nerve ring, which flows continuously across each arm-to-arm junction while also linking the neuropil of neighboring arms. The AAUT and inner nerve ring subdivisions were labeled to provide context to the segmentation shown in (A). B.) Higher-power examination of an arm-to-arm junction reveals microanatomical features of the octopus at the cellular level.

### Long range behavior-to-brain neural axes for chemotactile sensing, integration and memory

Long-range connections between sucker and brain were demonstrated by fine chemical and tactile sensing by suckers in behavioral experiments with live *O. bimaculoides* (Buresch et al., 2022, 2024; Sepela et al., 2025; van Giesen et al., 2020; Wells, 1978a; Wells & Young, 1969) and by loss of chemotactile learning and memory observed after ablation of the “inferior frontal system” (i.e., inferior frontal/subfrontal/buccal lobe complex) (Wells, 1978a). Past evidence of the axonal pathways connecting suckers to these brain lobes included silver-stained neurons in portions of multiple specimens (Young, 1971) (Young, 1971). The resolution of our detector system does not distinguish single small axons or nanoscale synaptic ultrastructure, but textures were sufficient to derive a context-sensitive understanding of the entire behavior-to-brain neural axis for chemotactile sensing, learning and memory. Such long-distance 3D chemotactile connectivity networks are illustrated in an oblique section through the scan (Figure 2B). The basic neural pathway is sucker → axial nerve cord of the arm → arm ganglion → nerve ring (to link additional arms) → brachial lobes of CNS → inferior frontal system and subvertical lobe of the CNS. Our continuous segmentation of this complex network includes elements from both the peripheral and central nervous systems, over which behavioral control appears distributed (Figure 2A).

To better understand how the sucker communicates with the brain, we characterized local connections between suckers in the hatchling and adult arm sample to the equivalent of each arm “spinal cord:” the axial nerve cord (ANC, Figure 3). Starting from the sucker, we traced meandering sucker (oral) nerves aborally, through muscle and neuronal cell body layers, into the neuropil of the ANC where they closely approach a pair of longitudinal tracts. These paired longitudinal tracts within the ANC neuropil span the length of each arm, and run bilaterally and parallel to the cerebro-brachial tract (CBT), the major nerve cord located along the aboral edge of the ANC. Their intermediate position between oral and aboral elements of the ANC gives rise to the name *intermediate longitudinal arm tracts* (iLT, Figure 3, Video S4 iLT). Based on 3D topology and position, these tracts appear analogous to the “bilateral longitudinal tracts’ recently described in histological sections of the arm (Olson and Ragsdale, *biorxiv*, 2025); our whole-animal volume reveals their continuous course and connectivity *in situ* (Olson & Ragsdale, 2025). We observed several projections from sucker nerves merging with these parallel tracts, revealing a longitudinal architecture that links the suckers of each arm via a pathway distinctly separate from the CBT.

Since neural and other structures of older animals are larger and more readily resolved, we also examined the 3D histotomogram of a distal portion of an excised adult arm (Figure 3B-E). There we could corroborate pathways that are less pronounced in the hatchling to better address questions about communication between suckers and brain raised by Graziadei and Wells (“Chapter 3: The Nervous System of The Arms,” 1971; Wells, 1978a; Young, 1971). The adult arm data allowed us to validate previously un-characterized chemotactile pathways, first seen in the hatching, that link sucker nerves in the arm to the brachial nerve and, in the complete hatchling scan, with multiple brain control centers involved in integrative sensory processing, learning and memory (“Chapter 3: The Nervous System of The Arms,” 1971; Wells, 1978a; Young, 1971).

We followed the longitudinal tracts in the adult arm fragment for long distances in the hatchling reconstruction, despite the smaller arm structures and artifacts associated with sample movement. Segmentation of the paired tracts forming the iLT within the proximal ANC neuropil revealed connections with the CBT as the cell body layer thins and disappears and the CBT becomes the brachial nerve at the proximal portion of the arm (Figure 3A). These new connections complete an anatomical path from sucker to central brain. Fascicles of the iLT appear [to maintain a compartmentalized bundle] within the oral / medial aspect of the brachial nerve as it projects towards the brain (Figure 3). While continuous delineation of bundle boundaries within the proximal brachial nerve is presently difficult, we hypothesize that axons in the iLTs defasciculate as a group upon entering the brachial lobe a subset contributes to cerebrobrachial connectives that lead towards the buccal and frontal lobes (Figure 2). Within the subesophageal brain, other brachial nerve axons then diverge into fascicles that form stereotyped, symmetric arches. We use *brachial arcade* (BrA) to denote these repeated bifurcations as they project to join the interchromatophore tracts, brachio-palliovisceral and cerebro-brachial connectives, and the brachial lobe itself (Video S5 BrA, S6 BrA Chromat). These arches provide the first high resolution 3D confirmation of the complex wiring architecture schematized by (Budelmann and Young, 1985), demonstrating how the brachial nerve physically sorts mixed sensory and motor signals upon entering the central brain (Budelmann & Young, 1985). We expect interbundle connections to be more readily characterized as imaging precision improves.

### Mediators of adjacent arm communication: Arm-to-Arm U-tracts (AAUT) in the nerve ring

To address the question of what microanatomy mediates inter-arm communication in the octopus, especially in response to recently reported stimulation of the interbrachial commissure (Chang and Hale 2023), we segmented the entire circumesophageal brain, or “nerve ring”(Chang & Hale, 2023). The nerve ring is a large and complex axon-rich structure that loops around the proximal end of all eight arms and contains large commissures passing between arms, alternating with smaller nerve bundles covering the proximal end of each arm (Videos S7 AAUT 3D) (Graziadei and Young, 1971) (Young, 1971). Serial examination of slices in planes approximately parallel with nerve pathways clearly revealed multiple fascicular elements within the nerve ring (Video S8 AAUT 2D). Some run from arm ANC to arm ANC, providing a distinct route for communication between adjacent arms. We call these U-shaped fascicular patterns, which are consistent across all eight commissures of the nerve ring (Figure S5), Arm-to-Arm U-Tracts (AAUTs) (Figure 4, Figures S5). These distinct tissue domains within the nerve ring (Figure S4; video S8 AAUT CT 2D) are corroborated by laser-scanning confocal microscopy (LSCM Video S9 AAUT LSCM 2D), and histology.

The AAUTs form a continuous neural highway between adjacent arms. Proximally, AAUT fibers ascend along the lateral margin of one arm’s ANC, arc across the interbrachial commissure, and then descend along the corresponding margin of the neighboring arm’s ANC. There, the AAUTs appear to converge with the brachial nerve to form the CBT. Crucially, the AAUT from the left (−1) and right (+1) neighbor do not appear to defasciculate. Instead, they persist as distinct lateral fascicles that continue down the length of the arm, bookending the central brachial nerve bundle (Figures 3B, 4). This architecture is consistent with a spatial arrangement of the nerve bundles that preserves spatial identity within each of the involved arms.

AAUT connections between neighboring arms through the nerve ring are similar between 3-days (Fig. S5D) and 3 months (Fig. S5B) of development. This geometry explains the short ≈24 ms electrophysiological time for signals to pass between adjacent arms (Chang and Hale 2023) (Chang & Hale, 2023). Furthermore, the absence of AAUT connections beyond immediate neighbors (−1/+1) provides a plausible structural explanation for the signal attenuation observed by Chang and Hale across distant arms. Overall, 3D rendering of the nerve ring and its segmented parts illustrates how whole-animal reconstruction at histological resolution allows visualization of previously less well characterized structures (see figures 3.11 and 3.12 in Graziadei and Young, 1971), including suggestions of roles the anatomical structures may play in coordinating prehensile behavior (Young, 1971).

### Democratized visualization of whole-organism microanatomy

Access to visualization tools is necessary to facilitate validation and discovery from whole-organism scans. The 2.6 TB size of the 16-bit centimeter-field submicron micro-CT reconstructions make it impractical for most workstations to take advantage of full-resolution, multiplanar 3D visualization. We therefore facilitate accessibility to interactive visualization in 3D using a standard web browser by having provided a modification of the software package Neuroglancer (Figure S8, Table S1), which includes the reconstruction and 3D labels(Maitin-Shepard et al., 2021). Neuroglancer provides a zoomable and rotatable 3D rendering of the reconstruction with and without labels, and 2D slice views in adjustable sets of 3 orthogonal planes, enabling exploration of the data in any desired orientation and magnification (Video S10 NG Demo, Maitin-Shepard et al., 2021)(Maitin-Shepard et al., 2021). Web links for each panel captured within Neuroglancer are provided within Table S1, enabling the reader to directly navigate to the exact corresponding location within the 3D volume. We provide a neurobiology-focused ontology that includes a hierarchical listing of the anatomical structures highlighted in this manuscript. All annotations, data, and segmentations are available for download on https://cephalopod.team/histotomography/Octo9/ [RRID: SCR_027759].

## DISCUSSION

We generated a first whole-organism 3D blueprint of *Octopus bimaculoides* at histology-like resolution using a novel wide-field micro-CT detector system. Within the resulting 2.6 TB digital volume we segmented every major organ system in a globally accessible web resource for community download, annotation and analyses. This whole-organism blueprint enables the study of long-range relationships and connections between organs and organ systems in the context of the whole animal. There remain in the data many intricate implicit anatomical relationships awaiting further explicit segmentation by the community. We focused primarily the chemotactile pathways as an exemplar.

Octopuses chemically and mechanically identify whatever they touch through chemotactile receptors on the surface of their suckers. Rapid neural integration of this information at multiple levels from periphery to CNS enables the animal to efficiently detect and interact with environmental objects (animal, plant, inert), and numerous studies have provided glimpses of parts of the circuitry underlying this ability(Neacsu & Crook, 2024; Olson, Moorjani, et al., 2025; Olson, Schulz, et al., 2025; Olson & Ragsdale, 2023). Graziadei and others described the organization and innervation of the many suckers studding each arm (“Chapter 3: The Nervous System of The Arms,” 1971; Neacsu & Crook, 2024; Olson, Schulz, et al., 2025; Olson & Ragsdale, 2023). Recent studies identified neural elements that suggest a basis for information exchange between neighboring suckers, yet they did not link this network to nerve tracts running centrally(Olson, Schulz, et al., 2025; Olson & Ragsdale, 2023). Microanatomical details of the nerve ring have been characterized in fragments for many years (Young, 1971)(Young, 1971). Likewise, extensions of the CBT (cerebro-brachial tract) projecting as brachial nerves into the CNS have long been recognized (Budelmann and Young, 1985), as have several central sources and targets - chief among them (for taste and touch) being the inferior frontal lobe (IFL) complex (Budelmann & Young, 1985). Ablation studies of this complex have demonstrated that the animal requires this central brain region to identify and integrate tactile and chemical information throughout the body (Wells, 1978a, 1978b; Wells & Young, 1969, 1972). The IFL shares this information with more executive lobes such as the subvertical lobe, thought to be the center for chemotactile learning and memory (Figure S9) (Abbott et al., 1995; Young, 1971). Together these historical observations constitute a plausible connectivity map for this sensorimotor system. Herein, we present the architecture of long-range pathways delivering chemotactile information to the brain within one specimen. By detailed study and segmentation of grey-scale textures in our microCT data, we defined a mesoscale connectome at the nerve-bundle level, including detailed 3D highways from the periphery to the central brain. Our evaluation process included identifying and segmenting previously uncharacterized nerve tracts linking oral nerves from the suckers to the brachial nerves and into the brain (ILTs, Figures 2 and 3). Our whole-body tracing places the local tract architecture recently described in the arm (Olson and Ragsdale, *biorxiv*, 2025) into a long-range context and reveals how these longitudinal systems interface with classic central targets such as the inferior frontal and buccal lobes (Olson & Ragsdale, 2025). Our work allowed us to trace connections within the brain from the brachial nerves through the cerebrobrachial connectives into the buccal and inferior frontal lobes. The structures are, for the first time, continuously visible within a single set of images spanning a single animal.

In addition to probing chemotactile wiring across the whole-octopus, our high-resolution volumetric datasets enabled spatial interrogation of inter-arm connections. Arm coordination by the octopus is well-studied for its implications across soft robotics, prosthetics, and neural network design, but little is known about the hierarchy of neuronal wiring that enables the animal’s semi-autonomous manipulation within and between its 8 muscular hydrostat arms. Recent electrophysiology studies have provided evidence for local signal transduction through the interbrachial commissure linking neighboring arms (Chang and Hale, 2023), and behavioral tracking has shown that octopuses preferentially recruit adjacent arms when foraging in visually occluded environments (Gendreau et al. 2025) (Chang & Hale, 2023; Gendreau et al., 2025). However, the specific structures that facilitate this communication remain incomplete (Chang & Hale, 2023).While longitudinal tracts have recently been identified within the arm neuropil, their continuity with the interbrachial commissure has remained unresolved (Olson and Ragsdale, *biorxiv*, 2025) (Olson & Ragsdale, 2025). Graziadei suggested a possible subdivision of the nerve ring, but no 3D structures delineating this arrangement had been characterized. We segmented the AAUT subdivision of the nerve ring within all 8 interbrachial commissures of our hatchling dataset. These subdivisions link the cerebrobrachial tract of each arm to each of its neighbors. Furthermore, Chang and Hale (Chang & Hale, 2023) observed that neural signals attenuate as they travel to distant arms. Our observation that AAUTs connect only adjacent arms suggests that inter-arm signaling occurs via sequential neighbor to neighbor transmission, likely necessitating attenuation-inducing relays to reach distant arms. Beyond signal transmission, this restricted topology offers a candidate structural solution to the motor control paradox described by Zullo et al. (2009) (Zullo et al., 2009). If the central brain issues generalized motor commands to multiple arms simultaneously, a peripheral mechanism is required to prevent limb interference. The AAUT provides the requisite architecture for neighbor-state signaling, ensuring necessarily distinct kinematic states during the execution of global motor programs (Zullo et al., 2009). Within the tomographic reconstruction of the excised adult arm, we identified these distal extensions of subdivisions as contralateral components of the CBT. We were unable to resolve the AAUT at the very distal aspects of the arms due to non-linear motion artifact and extension of some arms beyond the field of view. The AAUT subdivisions appear to exchange fibers both within the nerve ring and within the axial nerve cord of the arm – suggesting a potential mechanism for crosstalk between neighboring axial nerve cords and possibly the central brain.

An unresolved question in cephalopod neurobiology is how the octopus directs motor commands to a specific arm. Functional studies of the octopus brain demonstrated a non-somatotopic organization, where stimulation elicits distributed multi-arm responses rather than single-arm movements (Zullo et al., 2009).This suggests that the specificity of arm recruitment is defined by the complex wiring of the architecture of the peripheral nervous system rather than a central topographic map. The system might rely on peripheral sensory input to direct these central commands to specific arms. By providing a link from the suckers to the inferior frontal system, the brain’s primary chemotactile integrator, the iLT is anatomically poised to deliver the localized spatial cues required to gate these global motor programs to the appropriate limb. Neuroarchitecture evaluated at the mesoscale can inspire future electrophysiological experiments, such as directing where to place electrodes to query the function of the iLT.

This study represents an advance in microscopy and 3D histology that has specific limitations. The small sample number used in this study means that some of the described features may differ with anatomical variation. Motion artifact also limited the resolution of the reconstruction in certain areas (see movement correction section), which can be addressed by rigidly embedding the samples (Lin et al., 2018). Synchrotron X-ray beams are improving in flux and stability, which should improve signal to noise and shorten imaging time. The newly upgraded 2-BM beam height at Argonne National Laboratory can be modified to accommodate the full vertical FOV (7 mm) of our detector to lessen the need for multi-scan acquisitions. The broader application of centimeter-scale histotomography such as sets of mutants and chemically-treated animals and plants, and to tissues from larger organisms such as humans would be greatly facilitated by the creation of synchrotron beamlines dedicated to biology.

Overcoming these imaging barriers is the first step toward a larger goal. Ultimately, *digital organismal biology* seeks to extend organismal biology by pairing open, navigable whole organism digital specimens with models that connect structure to physiology, behavior and environmental drivers. Our octopus dataset provides one such anatomical basemap: histology-like detail across an intact organism, preserved *in situ*, on which functional measurements and hypotheses can be registered, queried and tested *in silico*, complementing lab and field work. The same workflow generalizes to other centimeter-scale organisms (e.g., fish, arthropods, plants) and tissue samples from larger organisms, enabling cross-taxon comparisons of organ systems and long-range tracts. We frame the present work as an anatomical foundation for *digital organismal biology*. Functional integration can follow via data registration, modeling and targeted validation and potentially by creation of various tracers and biomarkers detectable by X-ray (Katz et al., 2021). In sum, democratization of imaging and resulting 3D datasets is intended to foster collaborative segmentation and other methods of analysis and may serve as a scaffold for hypothesis generation and future interdisciplinary science.

## Supporting information

Video S1 Tour

Video S2 oINC 3D

Video S4 iLT 3D

Video S5 Brachial Arcade

Video S6 Brachial Arcade Chromat

Video S7 AAUT 3D

Video S8 AAUT CT 2D

Video S9 AAUT LSCM 2D

Video S10 NG Demo

## ACKNOWLEDGEMENTS

We dedicate this work to the memory of Cheng lab team members Damian Van Rossum and Peggy Hubley, who played key roles in making this work possible. We thank Pavel Shevchenko, Viktor Nikitin, Alan Kastengren, Alex Deriy, and Francesco De Carlo of Argonne National Labs for their support and valuable insight in conducting synchrotron micro-CT experiments, Phil Vargas for discussion of micro-CT reconstruction optimization, Marianne Klinger of the Penn State Pathology Core for octopus histology and H&E staining, and Jean Copper for helping with scanning of octopus slides through the Penn State Pathology Slide Scanning Core. Imaging work was made possible by funding from NIH grants 1R24OD18559, R24OD035407 and from the Jake Gittlen Laboratories for Cancer Research and Penn State Cancer Institute and to KCC. RTH and SLS gratefully acknowledge support from ONR Grant N00014-22-1-2208. We acknowledge the Central Microscopy Facility (CMF) at the Marine Biology Laboratory for the use of their confocal microscope. We acknowledge the involvement of Zebrafish Functional Genomics Facility (RRID:SCR_021199) in the development of histotomography and instrumentation. This work utilized cost-free academic use of Dragonfly software (Object Research Systems, Montreal, Canada) for image analysis and visualization and thank the support team at Object Research Systems (ORS), especially Hubert Taleb, for assistance with its use. We are all indebted to three pioneers whose meticulous research on octopus neuroanatomy inspired and enabled this report: J.Z. Young, Martin Wells and Pasquele Graziadei.

## Materials and Methods

### Tissue Preparation

Octopus bimaculoides hatchlings were obtained from eggs laid by a female octopus captured in the wild (southern California; Aquatic Research Consultants, San Pedro, CA 90732) and sent to Woods Hole. The female was housed and fed according to well established protocols at the Marine Resources Center of the Marine Biological Laboratory. The female was normal and healthy in all respects, and she brooded her eggs in typical fashion until they hatched after full development (Gendreau et al., 2025). Since O. bimaculoides has no paralarval or metamorphic phase, their hatchlings are fully formed “miniature adults.” that possess the anatomy and behavioral capabilities of juveniles and adults (aside from reproduction). For tissue preparation, hatchlings were anesthetized in sea water containing several percent ethanol and/or magnesium chloride, then placed in a fixative of 4% paraformaldehyde in sea water for several days (van Giesen et al., 2020). One 3-day old hatchling was immersed for a week in several % phosphotungstic acid and eosin [AS1].

The tissue was rinsed several times in distilled water, then chemically dehydrated with acidified 2-2-dimethoxy propane (DMP) overnight in a closed container. To reduce surface tension while drying, it was immersed in the low vapor-pressure liquid hexa-methyl di-silazane [AS2] (HDMS) in a partially covered petri dish and was left undisturbed in a hood for several hours until the liquid had fully evaporated. The intact hatchling animal (“prep #MBL9”) was gently glued to the surface of an aluminum SEM stub mounted with plasticine onto a ThorLabs ½” diameter optical post [TR1] screwed into a 1” x 1” kinematic base [KBT1X1]. To minimize attenuation of the illuminating X-ray beam, these preps were not embedded (e.g. in epon [AS3], as commonly used for TEM). For adult arm imaging, a short (∼1/2 inch long) distal section of arm L2 from a euthanized adult Octopus bimaculoides (prep #MBL7) was fixed in 2% glutaraldehyde with 2% paraformaldehyde for 24 hours at 4 degrees C, post-fixed in 2% osmium (on ice) for 15 minutes, then stained with ∼0.5 % PTA (2 days, RT), and 5% strontium chloride (3 weeks, RT), both in 70% ethanol. The arm was dried and mounted similarly to the MBL9 prep.

### Histopathology

Histological processing of octopus tissue was performed using standard paraffin-based methods. Whole specimens were fixed in 4% paraformaldehyde in phosphate-buffered saline (PBS) to preserve tissue morphology, followed by graded ethanol dehydration and paraffin embedding. Following fixation, tissues were dehydrated through a graded ethanol series (70%, 80%, 95%, and 100%) and cleared in xylene (Fisher Scientific, Cat. #X3P), then infiltrated with and embedded in paraffin (Paraplast Plus; Leica Biosystems, Cat. #39601006). Serial sections were prepared in the dome-to-arm orientation at a thickness of 5 µm using a microtome and were mounted onto Superfrost Plus slides (Fisher Scientific, Cat. #12-550-15). Odd number sections were stained with hematoxylin and eosin (H&E) to visualize cellular and tissue architecture. Even number sections were left unstained, to be used in the future. Coverslips were applied using mounting medium (Permount; Fisher Scientific, Cat. #SP15-100). Stained slides were digitized with a whole-slide brightfield scanner (Aperio AT2, Leica Biosystems) at 40× magnification, producing high-resolution images for visualization and digital reconstruction of serial sections.

### Synchrotron Imaging

Histotomographic image acquisition was performed at the sector 2 bending magnet beamline (2-BM) of the Advanced Photon Source (APS) at Argonne National Laboratory. To align to the beam, two Newport MTN100PPs were used for horizontal and vertical movement and IDL280 for vertical movement. The sample to scintillator distance (SSD) was 70mm. The QRM phantom (Fig. S3) and the octopus (Fig. 2) were illuminated with monochromatic 25.5 keV light. Resolution of the QRM phantom scan (Fig. S3) was calculated by taking a 10-slice z-projection (average intensity) of the L feature in the reconstructed *tif stack. The edge spread function (ESF) was then taken from the long side of the L using the rectangular selection tool and plot profile function in FIJI. Bicubic spline interpolation was used to fit a curve to these sampled points, and the derivative of this curve was taken as the line spread function (LSF). The full-width half-maximum of this curve was then taken to be the empirical spatial resolution of our system.

Our system’s optics were designed to image the full height of the sample in a single visual field. Since 2-BM’s beam is less than 1 cm tall, 2 separate scans were utilized to cover the sample’s full height (Fig. S2). These scans were digitally reconstructed and combined to form a 3D volume within which we were able to identify every tissue type and organ system. To facilitate visualization and study, we utilized 3D viewers including Dragonfly and Neuroglancer (Maitin-Shepard et al., 2021). Cells larger than about 1.5 microns in diameter, groups of axons, large vessels, and ducts were readily discernable, especially in serial digital slices. Labeling components of the 3D structures (“segmentation”) derived from every major organ system. Whole-organism blueprints (Figure 1) provide context for understanding how microanatomical components of each organ system fit within the whole of the organ system and the whole-organism.

Tomograms of the octopus were acquired across 360 degrees and 9001 projections with a 28% projection overlap to minimize artifacts. These projections were then reconstructed into a 3D image volume using the gridrec algorithm from TomoPy (Gürsoy et al., 2014). A projection from angle 360 was registered to angle 0 for offsets in x and y to account for sample dropping 20 microns in y and shifting 2 microns in x during the scan. Projections were then linearly shifted in x and y to account for the offsets recorded, as a function of angle theta. Stripes were removed using the Fourier-Wavelet based method with a db25 and sigma of 2.4, a bilateral filter in the projection domain d,sc,ss = [3,1,1] [opencv], and BAC filter of alpha = 2.0, gamma = 2.0 were incorporated during the reconstruction step (Gürsoy et al., 2014; Witte et al., 2009). A subset of reconstructed volumes from each section were registered using the pairwise-stitching plugin in imageJ (Preibisch et al 2009) to record x y and z discrepancies between the two. Scans were then linearly combined based on these offsets (Preibisch et al., 2009; Schindelin et al., 2012, 2015). To measure the resolving power of our detector system across the full 10 mm FOV, we imaged a QRM Nanobar phantom (Cheng et al., 2011; Schindelin et al., 2015; Yakovlev et al., 2022). The effective pixel size of our detector system is 0.7 x 0.7 micrometers after magnification by the lens objective. As a measure of spatial resolution, the Line Spread Function (LSF) was calculated using an edge of the L feature of the phantom (Figure S3.B’L) yielding a Full-Width-at-Half-Maximum (FWHM) of ∼1.6 micrometers in the projection domain and ∼2.3 micrometers in the reconstruction domain (Figure S3.E).

### Segmentation of Octopus Anatomy

Segmentation of anatomical structures from the reconstructed micro-CT volume was performed using Dragonfly software (Object Research Systems; RRID:SCR_025150). To manage the large dataset (2.6TB) efficiently, regions of interest (ROIs) containing specific structures, such as the brachial nerves, nerve ring, and intramuscular nerve cords (INCs), were excised as cropped regions (crops). These crops were then loaded into Dragonfly for interactive segmentation.

### Sparse Labeling and Interpolation

We employed a sparse labeling approach combined with morphological interpolation to delineate target structures. This method was chosen due to its efficiency in handling large datasets and its suitability for segmenting elongated, continuous structures. Structures were sparsely labeled at regular intervals along a single axis, typically the axis exhibiting the clearest anatomical definition. Dragonfly’s morphological interpolation tool was then used to propagate these sparse labels along the chosen axis, generating a continuous segmentation.

### Path-Based Resampling for Large Structures

For elongated structures that spanned volumes too large to be handled in RAM, we created a custom path-based resampling approach. We first annotated the trajectory of the large structure using Neuroglancer by placing a series of points along its path (Maitin-Shepard et al., 2021). These annotations were interpolated to generate a continuous centerline representing the structure’s course through the volume. Using this centerline, we resampled the volume along the path of the structure, effectively straightening the curved structure into a flattened representation. This resampling reduced the data size by aligning the volume with the structure’s orientation, allowing us to work with a smaller, more manageable dataset. Segmentation was performed on this resampled volume using sparse labeling and interpolation. After segmentation, the labeled data were mapped back into the original volumetric space through an inverse transformation, restoring the segmentation to its correct anatomical location within the full dataset.

### Organ and Brain Segmentation

To segment full organ systems, the volume was resampled to 5.6µm resolution to allow access to the entire dataset in memory in Dragonfly (Object Research Systems; RRID:SCR_025150). Segmentation of large organ systems spanning across the organism and the initial segmentation of the brain lobes were performed on this volume via sparse labeling, interpolation, and smoothing. Intensity-based thresholding was used to remove empty areas around complex structures such as the gills. Brain lobes were identified based on neural texture, white and grey matter, position and connectivity with reference to existing literature, primarily (Bennice et al., 2025; Budelmann & Young, 1985; Young, 1971). After the initial pass at 5.6µm resolution, the brain lobe segmentations were resampled to the full 0.7µm voxel dimension resolution on a cropped brain lobe volume and refined with the smoothing and dilation tools in Dragonfly.

### Segmentation Validation and Processing

Segmentation accuracy from the nerve ring was assessed by comparing the 3D renderings of segmented structures with corresponding views obtained from confocal microscopy (see Confocal Microscopy Cross-validation) and in other cases by histology or serial slice visualizations. Following interpolation, the resulting segmentations were saved as grayscale files. To create a comprehensive representation of the segmented anatomy across the entire 2.6TB scan, individual cropped segmentations were combined into a compressed master file representing the geometry of the full 2.6TB scan. This master file series serves as a streamlined resource for visualizing and analyzing the 3D organization of the octopus nervous system.

### Confocal Microscopy Cross-validation

Confocal microscopy imaging was performed using a Zeiss 780 confocal laser scanning microscope with a 20x 0.8NA lens set to 0.6X zoom. The octopus was stained with Lucifer Yellow, carboxyfluorescein, and wheat germ agglutinin 633. It was fixed in 4% paraformaldehyde for 3 weeks, then sectioned freehand with a razor blade. Tissue sections were rinsed in distilled water then dehydrated with dimethoxypropane, cleared with xylene, and mounted between coverslips in Histomount. Each tissue section was imaged with 3 channels including 405nm, 444nm, and 594nm with ∼12% laser power.

## Supplementary Materials

### Supplemental Figures

**Figure S1:**
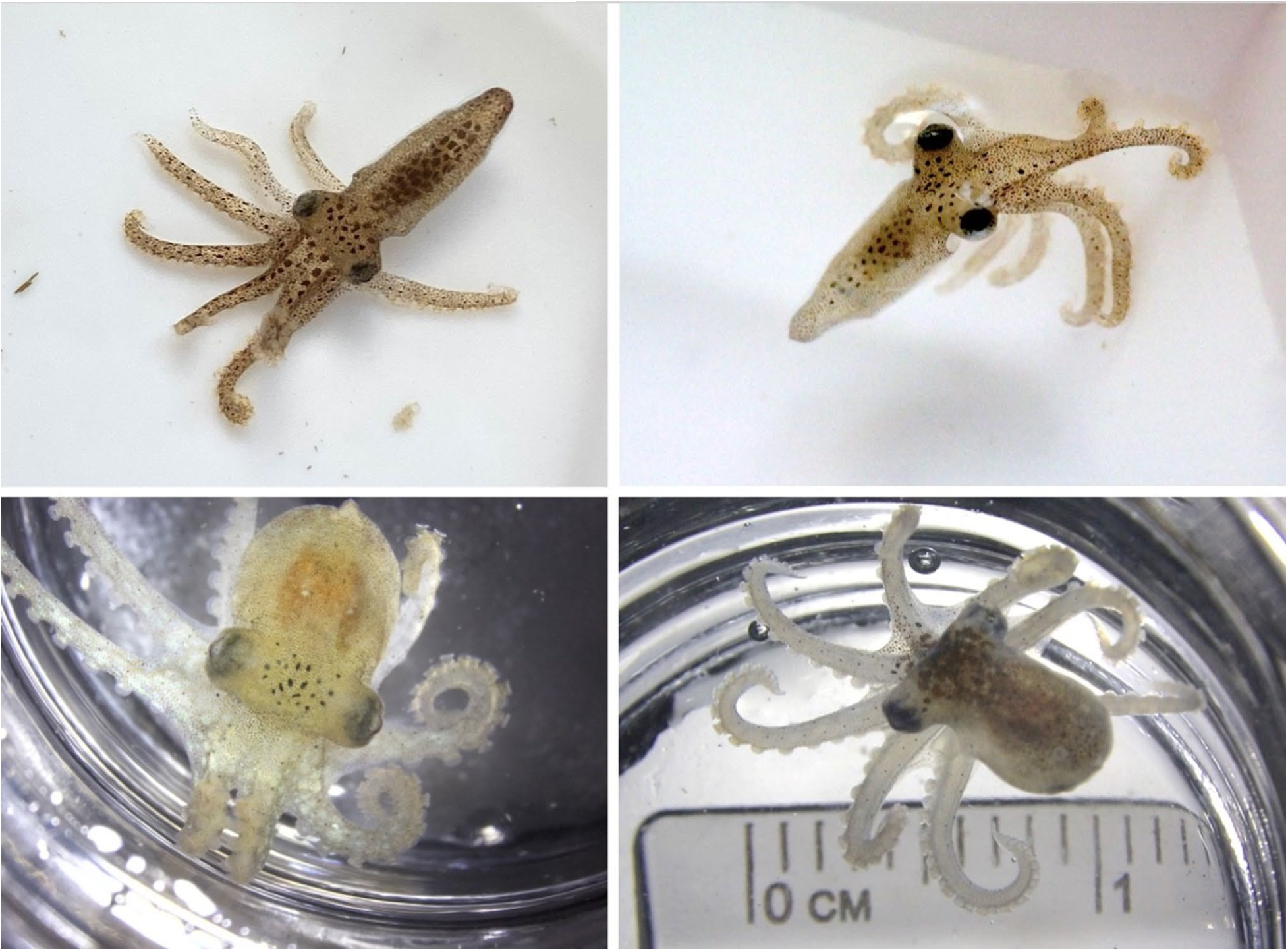
Hatchling *Octopus bimaculoides* are fully formed miniature octopuses. There is no larval or metamorphic stage; they become a typical benthic octopus immediately after hatching out of the egg. Mantle length is ca. 6-8mm.

**Figure S2:**
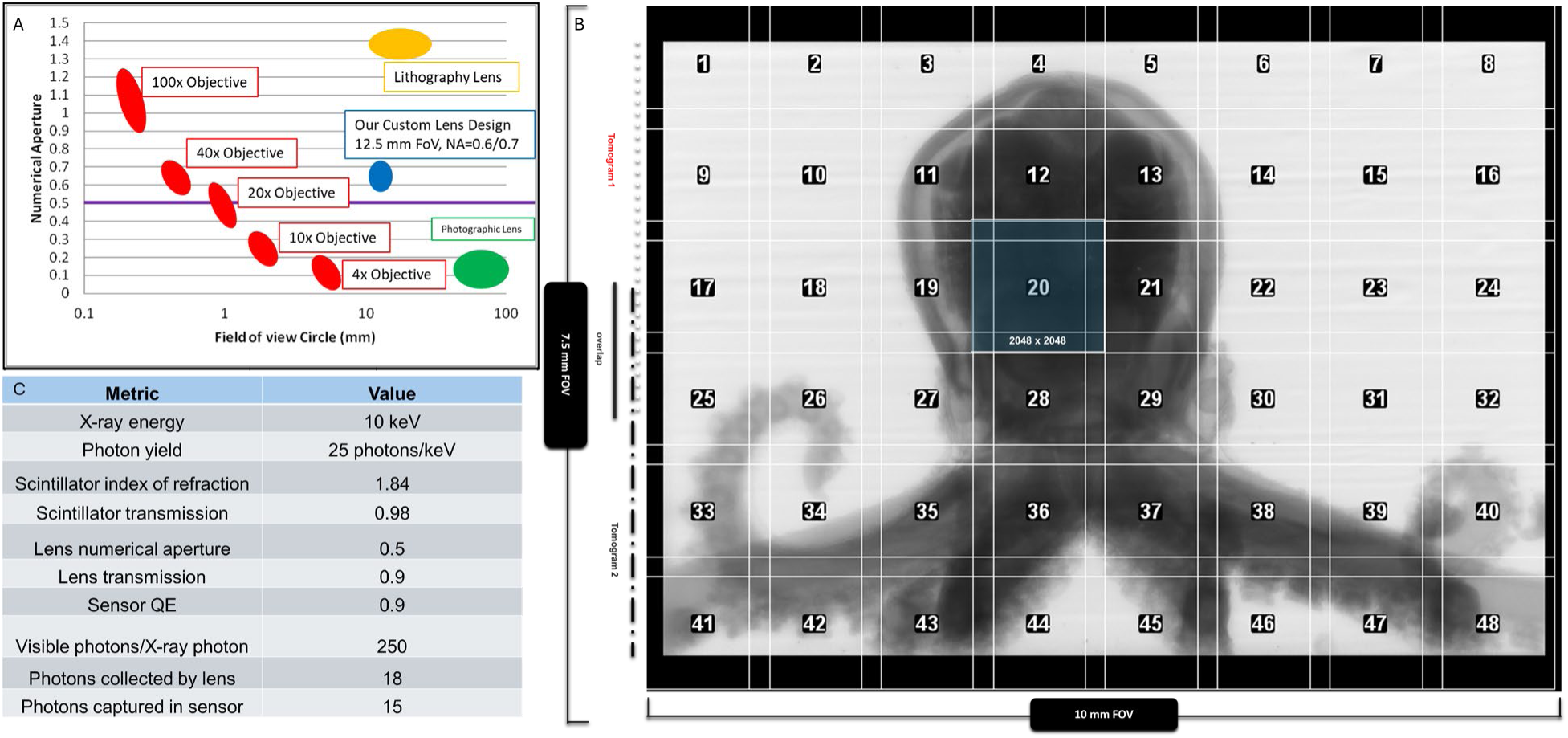
Contextualization of the 10 mm detector systems in relation to other imaging modalities. (A) Comparison of objective lenses by numerical aperture (NA) and FOV. Our custom lens is labeled in blue according to its diagonal FOV and NA. (B) 2 histotomograms of our hatchling octopus sample stitched in the projection domain with a grid overlay corresponding to equivalent area captured by a conventional 2k x 2k detector. The grid shows how many tomograms (48) would need to be computationally stitched at 15% overlap to cover the same FOV as our custom detector system. (C) X-ray sensitivity specifications for the sCMOS sensor employed within our custom system.

**Figure S3:**
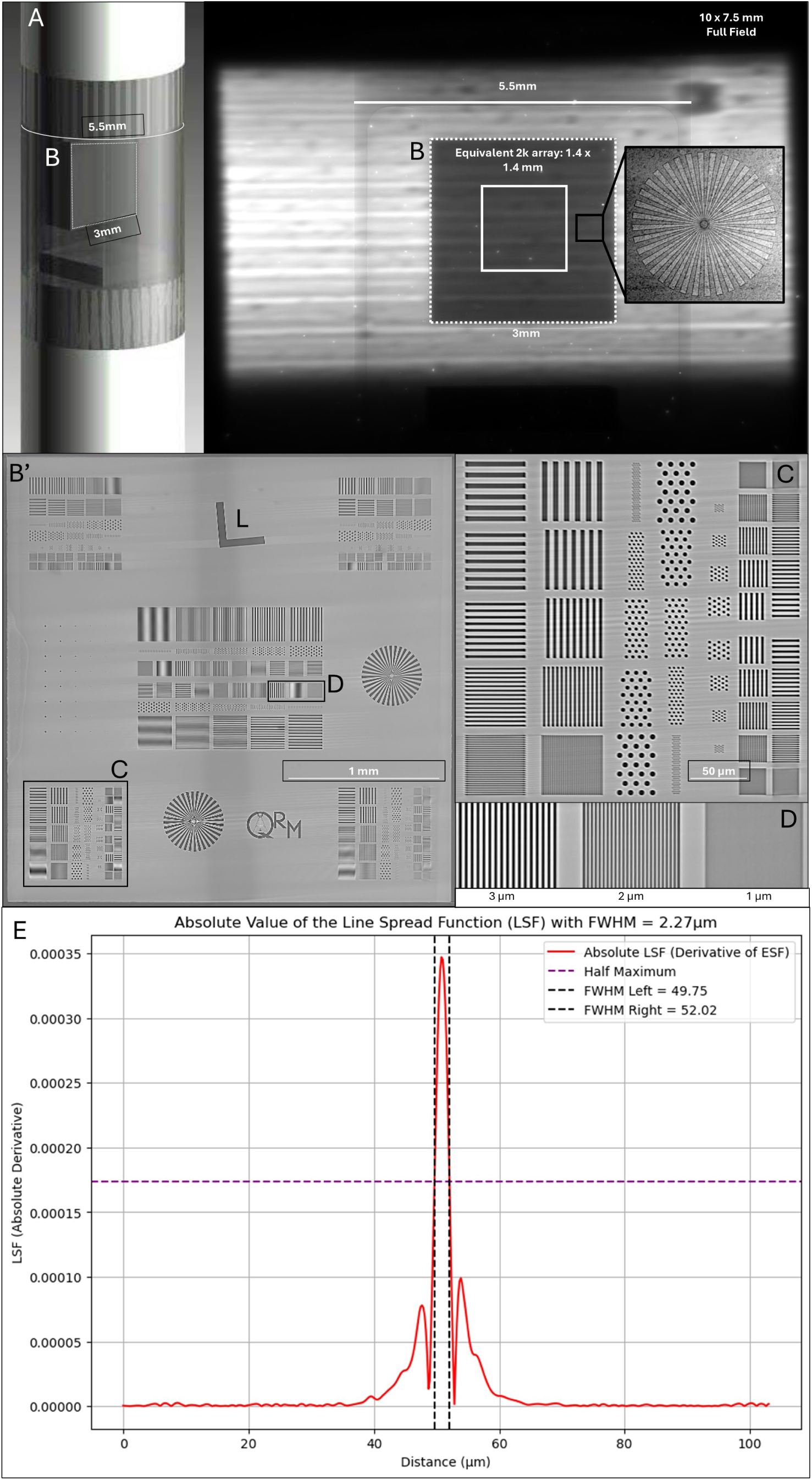
FOV and resolution characterized using a QRM phantom. A. 3D render of the phantom wafer embedded within its sample tube labelled for size. B. QRM phantom as seen in the projection domain prior to the initiation of a histotomographic scan. The full 10 x 7.5 mm field encompasses all surface area of the 5mm-wide sample tube as well as the full incident X-ray wavefront. The area that would be covered by a 2k x 2k pixel detector array of equivalent pixel size is represented by the solid white box within the phantom wafer (dotted white box). B’. Ten-slice average of the reconstructed QRM phantom showcasing the resolving power of our detector system. C-D. Higher-powered visualizations of the phantom line pairs and dot patterns in the reconstruction domain. E. Modulation transfer function (MTF) calculated from the L feature of the reconstructed QRM phantom. The MTF was taken as the discrete Fourier transform of the derivative of the edge-spread function recorded from a 10-slice average of the L visible in B’.

**Figure S4:**
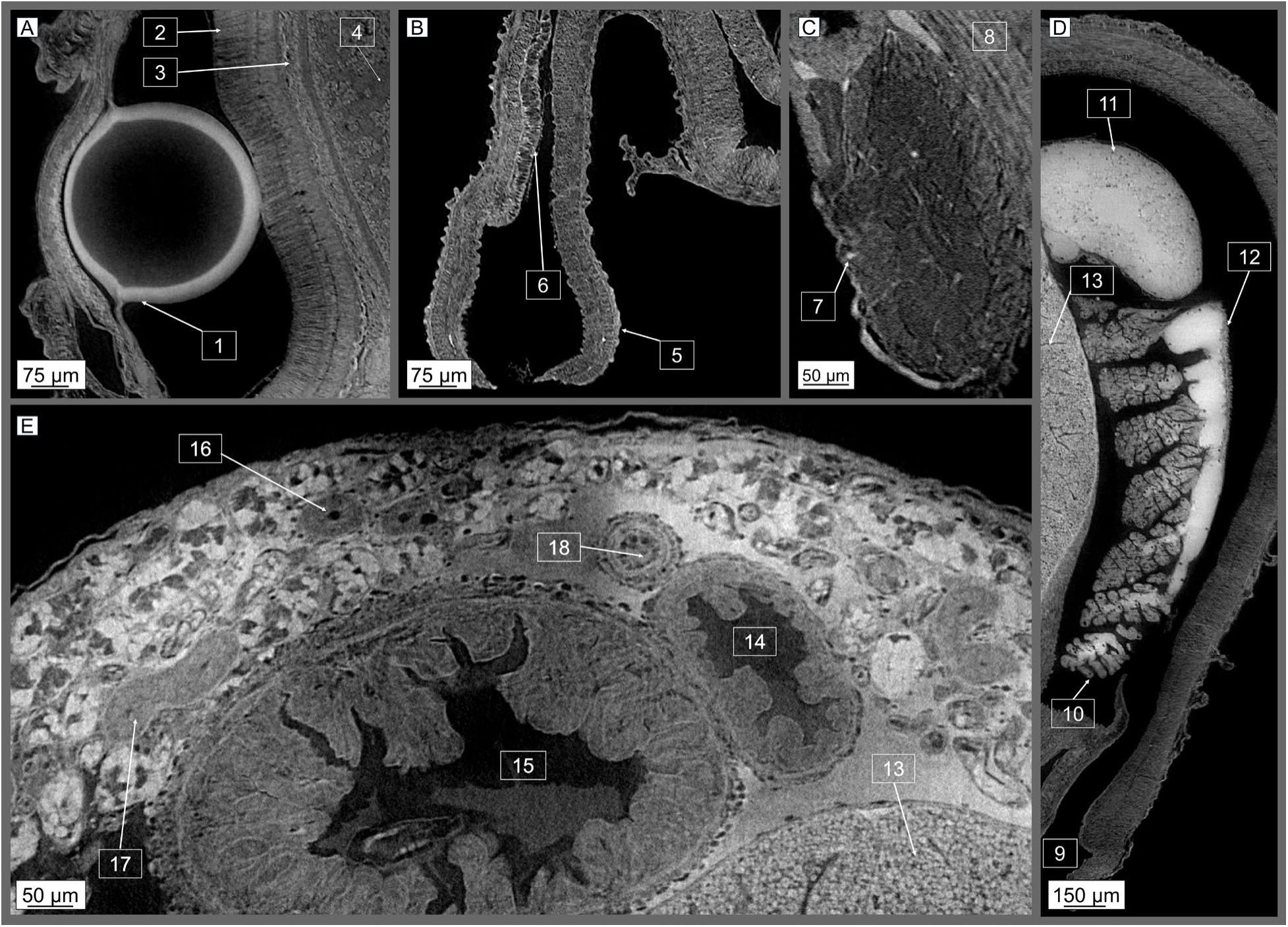
Whole-organism phenotyping allows for identifications of complex structures at the cellular level across multiple organ systems within a single scan. A: The Octopus visual system focuses light through the lens (1) and into a retina formed from long, single-type photoreceptor cells (2). Signals pass into the large, paired optic lobes, divided into the outer deep retina (3) and inner medulla (4). B: The octopus siphon (5) is used for respiration and locomotion. Glandular epithelium (6) lines a portion of the inner surface. C: The Stellate ganglia (7) control mantle musculature (8) through a series of radial stellar nerves. D: During inspiration, water travels through an opening in the collar (9) and proceeds across the gills (10), driven by muscle movements controlled by the stellate ganglions. The branchial hearts (11) perfuse the gills, separate to the systemic heart (not pictured), while blood departs through the efferent branchial vessel (12). E: A virtual slice shows parts of the digestive system, including the edge of the large digestive gland (13), esophagus (14), crop (15), and portions of the posterior salivary (venom) glands including glandular cells (16) and striated ducts (17). A major artery carrying blood from the systemic heart to the brain is also visible (18).

**Figure S5:**
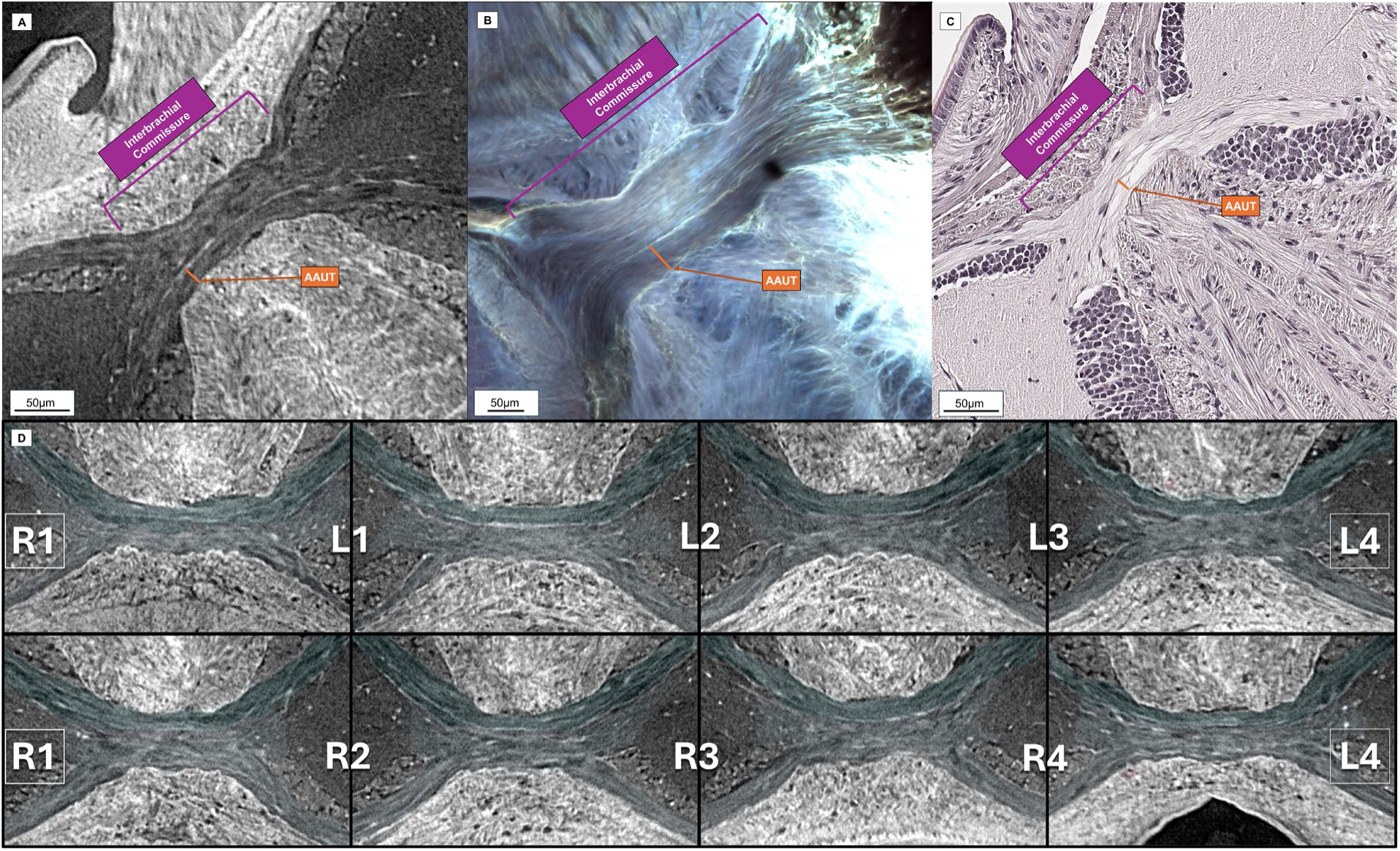
Axial cross-section of the interbrachial commissure as observed via histotomography (<u>left</u>) and confocal microscopy (right) Segmentations were hidden from the histotomography image for an unbiased visualization of the regional microanatomy. The boundary delineating the AAUT subdivision of the nerve ring appears as a white stripe in both imaging modalities. The histotomography image (<u>A</u>) was taken from a hatchling octopus, while the confocal image (B) was taken from a 3-month-old octopus. The axon bundles of the U-ring are labeled with orange bars in each image. (C.) Representative hematoxylin and eosin-stained section taken from the Interbrachial commissure of a separate hatchling octopus. (D.) Whole-animal imaging allowed us to create high-power 2D snap shots from each of the 8 interbrachial commissures including annotation of the corresponding segmented AAUT (teal green) and the inner nerve ring (blue). Each snapshot is oriented such that the top of the image is lateral to the octopus and the bottom of the image is medial.

**Figure S6:**
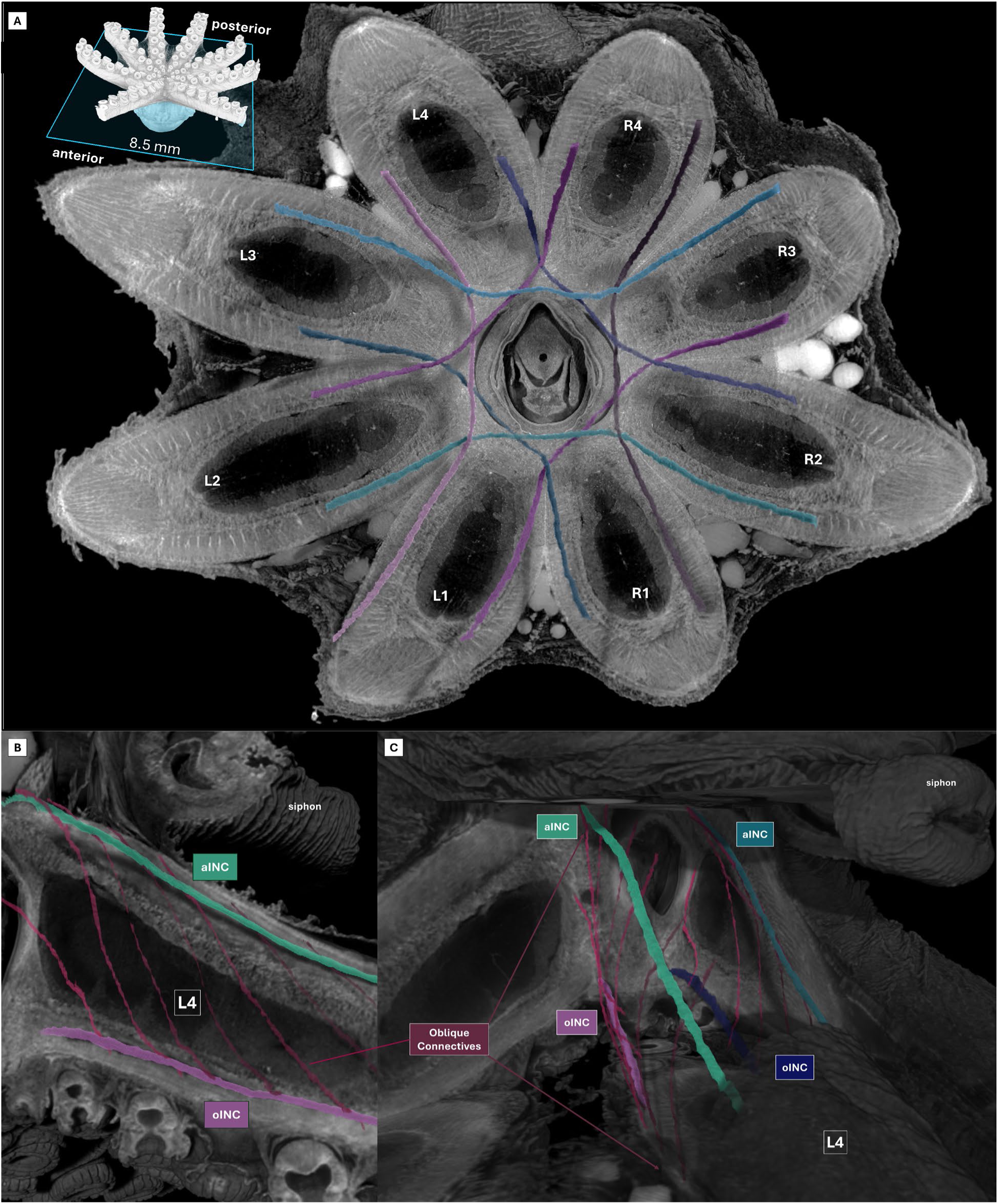
Wide-field histotomography enables continuous volumetric segmentation of the oral intramuscular nerve cords (oINCs) and oblique connectives (OCs) (<u>A</u>.) Transverse dig-ital cross section of the reconstructed octopus arm with 3D segmentations of the oINCs rendered in situ. (B.) Sagittal view of octopus arm L4 with rendered segmentations of ipsilateral oral and aboral INCs and their oblique connectives (OCs) that form an arcade-like motif. (C.) 3D rendering with cut out illustrating the anatomical context of the L4 OC within the broader volume of the animal. (C.) Full 3D segmentations of the oral INCs illustrating their overlap and tortuosity. R1 = Right arm 1, R2 = Right arm 2.

**Figure S7:**
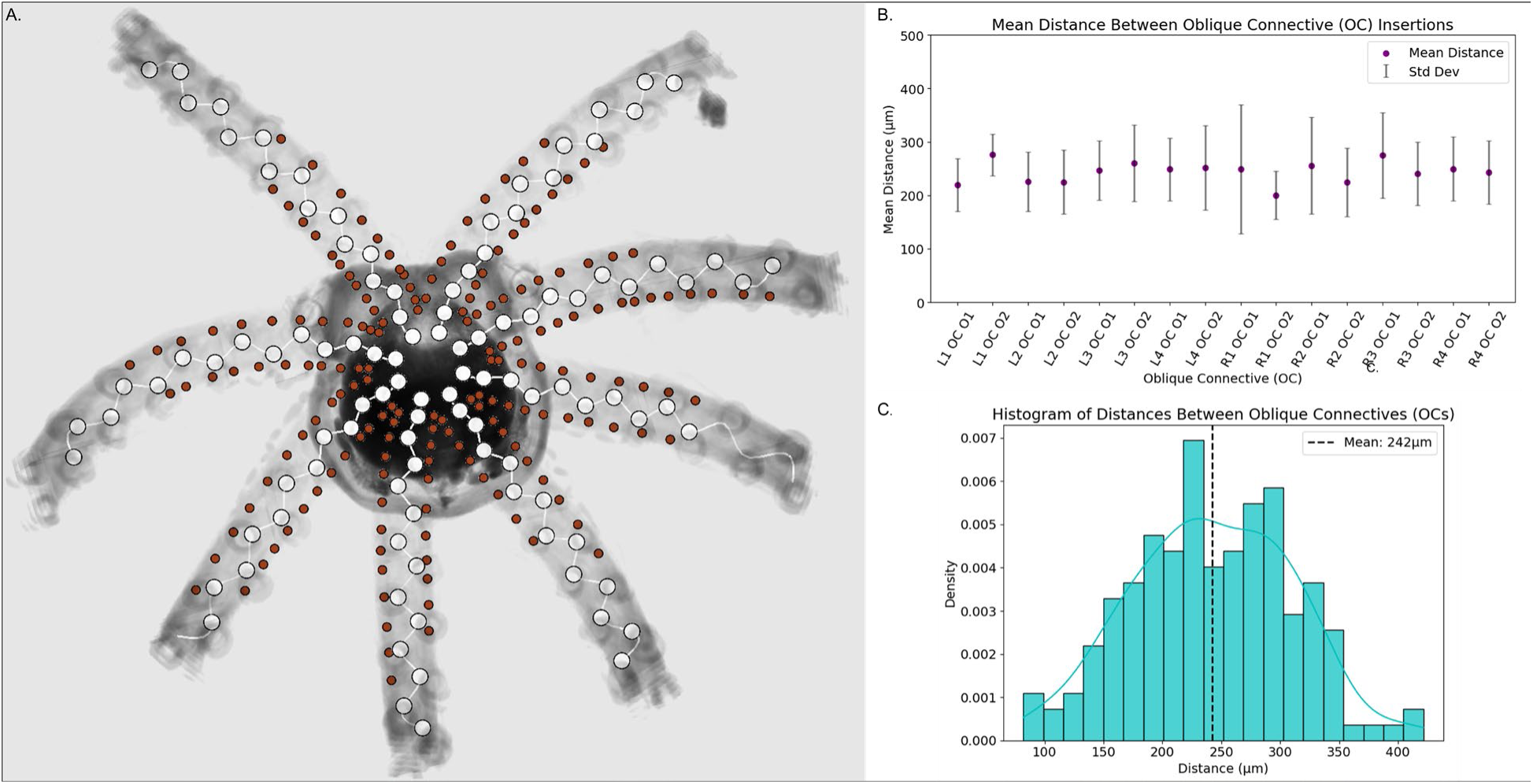
Euclidean distance between oblique connective insertions at the oral INCs and spacing of sucker ganglia reveal a similar pattern. A.) 3D point tracing of each visible sucker ganglion (white) rendered alongside point tracings of the oblique connectives (maroon) as they intersect with the oral INCs. B.) Mean and standard deviation of the Euclidean distance between the insertions of oblique connectives (OC) at each oral INC. Distances are plotted for each set of OCs, yielding 2 sets per arm (L1 OC O1, L1 OC O2, etc). C.) Histogram of all measured distances between the OC insertions not stratified by arm. The overall mean is indicated with a dashed line.

**Figure S8:**
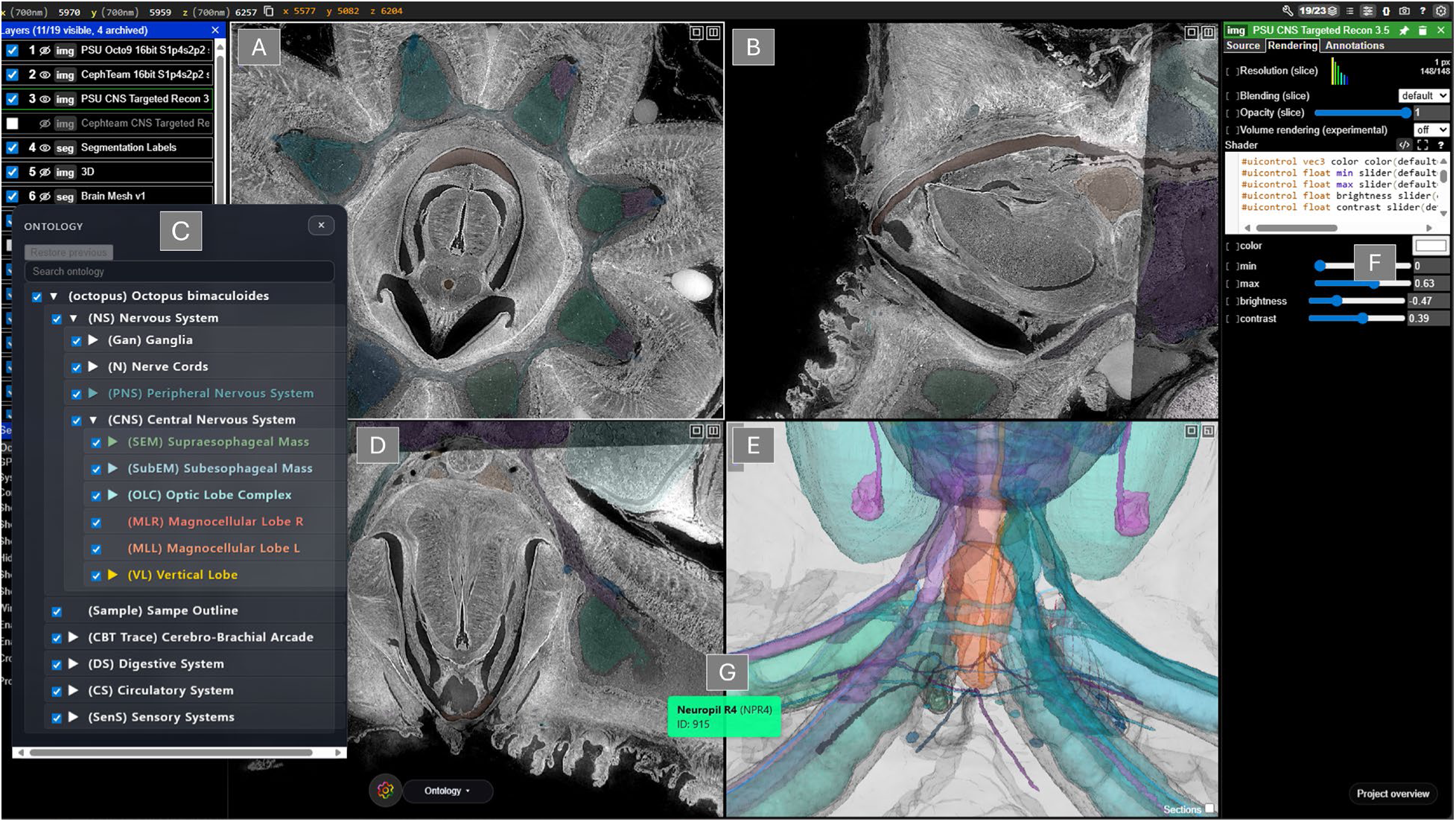
Quick start guide to navigate the image and segmentation using our customized Neuroglancer web interface. A) Transverse planar view of the octopus. Scrolling up parses distally while scrolling down parses proximally. B) Coronal planar view of the octopus. C.) The control panel on the left of the web page allows the user to select which elements of the dataset are visible, allowing them to hide segmentations, ontological labels*, 3D rendering, and the mi-cro-CT reconstruction itself. D.) Sagittal planar view of the octopus. E) Panel dedicated to 3D rendering of the reconstructed dataset and segmentations. F) Source, Rendering, and annotation control panel. The user can use this panel to not only alter the appearance of the image, but also to record coordinates of regions of interest and add their own custom point annotations and measurements by selecting “Annotations” at the top right and control-clicking on a voxel within panels A, B, or E. G) Example ontological label as seen within the web viewer. See Movie S9 for basic demonstration.

**Figure S9:**
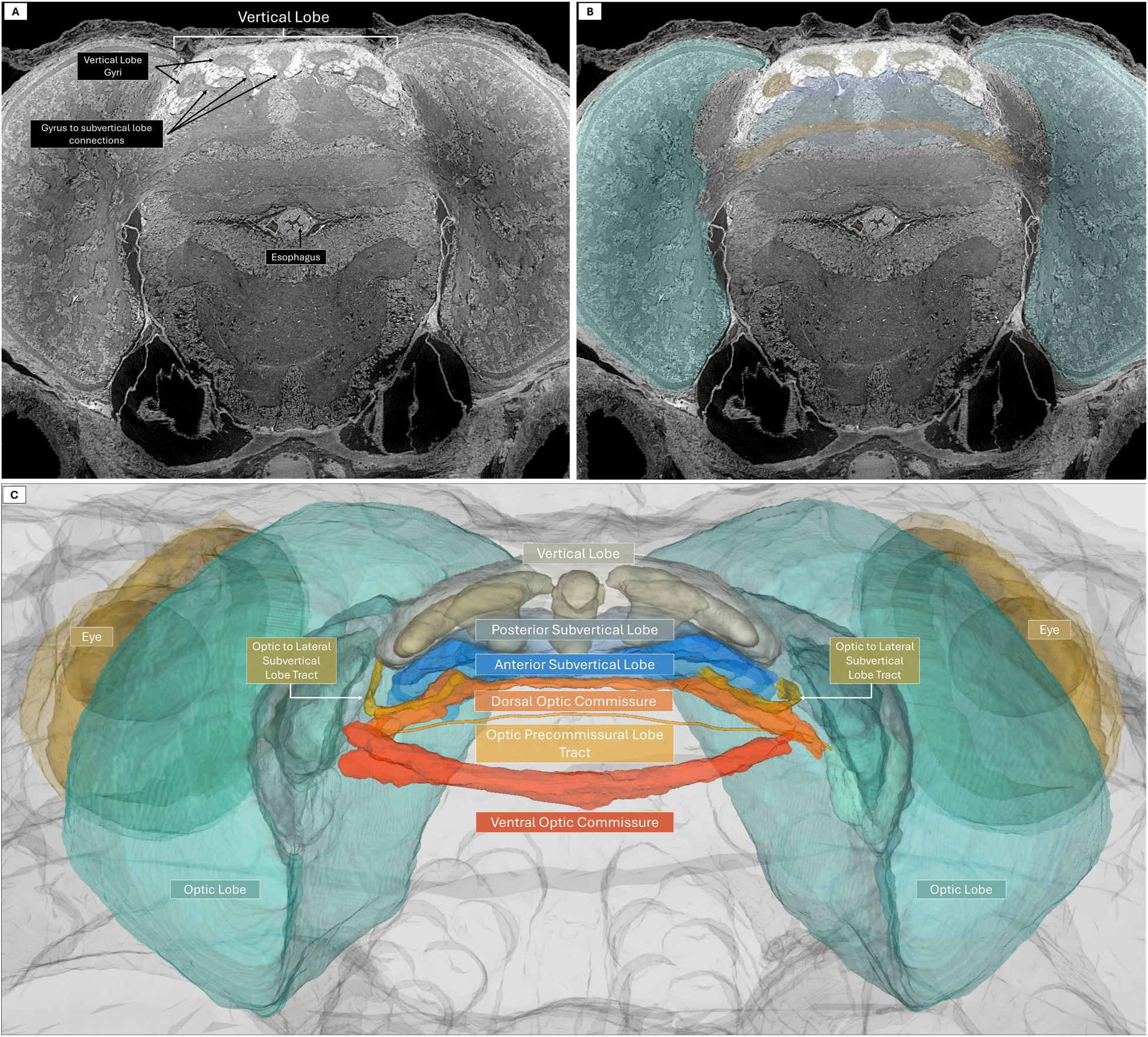
Major integrative connections to and from the subvertical lobe. A) Multiplanar visualization allowed validating alternative views of connections between the subvertical lobe and other brain regions that may play an important role in sensorimotor integration involved in learning and memory. A) Transverse 2D cross-sectional slices at 0.7-micrometer resolution used to recognize and color-label major connections in 2D, and example of which is shown in (B). Collective 2D segmentations validated by multiplanar inspection were used to create 3D representations shown in (C) that were then incorporated into the digital octopus. The subvertical lobes have extensive connections to other brain regions, the largest of which are the optic to lateral subvertical lobe tracts that link them to the larges brain lobes, the optic lobes after passing through the lateral subvertical lobes. C) The optic lobes also have direct connections to each other, primarily the large dorsal and ventral optic commissures, with some smaller paths such as the tract passing through the precommissural lobe. The eyes are lateral to the optic lobes.

### Videos

**Video S1: Tour of Octopus Histotomography (associated with Figure 1)**

**Video S2: 3D rendering of intramuscular nerve cord (INC) segmentations enabled by histotomography (associated with Figure S6).** The oINCs are rendered as colored 3D segmentations, illustrating their connectivity across all 8 arms.

**Video S3: 3D rendering oblique connective (OC) segmentations enabled by histotomography (associated with Figure S6).** The OCs are visualized as 3D segmentations and shown at multiple angles within the broader anatomical context of the arm.

**Video S4: Intermediate longitudinal tracts (iLTs) link oral nerves to the brachial nerve (associated with Fig 3)**. Oral nerve traces and segmentations are shown extending from the sucker epithelium through the ANC to the iLTs. Two segmented oral nerves (gray-purple hues) are shown with one projecting to the sucker ganglion. Segmented and traced oral nerves are shown touching the iLT in the spatial context of the peripheral neural architecture rendered *in situ* within the arm.

**Video S5: Brachial Arcade.** Neuroglancer rotational rendering of the Brachial Arcade (BrA).

**Video S6: Brachial Arcade and Chromatophore tracts.** Neuroglancer rotational rendering of the Brachial Arcade (BrA) and chromatophore tracts.

**Video S7: 3D segmentation of the Octopus nerve ring and AAUT in anatomical context (associated with Fig. 4**). The nerve ring volume of interest is shown in context of the full octopus reconstruction. The AAUT is a partition of the nerve ring that links brachial nerves of adjacent arms. Nerve ring segmentation is rendered in light blue, and AAUT segmentation is rendered in teal.

**Video S8: Slice-by-slice 2D view of nerve ring and AAUT segmentations within the histotomographic reconstruction (associated with Figure 4a**). Higher-powered view of the interbrachial commissure as visualized by histotomography reveals subdivisions of the nerve ring that define the AAUT. This video scrolls down and up through axial slices of the octopus while toggling segmentation masks of the nerve ring and AAUT to illustrate for the viewer how they were partitioned.

**Video S9: Corresponding slice-by-slice 2D view of nerve ring and AAUT as visualized by laser-scanning confocal microscopy (LSCM) (associated with Figure 4b**). This video highlights the same view as video S3, this time with confocal microscopy instead of histotomography. Here, the viewer is shown the stack of confocal images used for cross-validation in the segmentation of the AAUT.

**Video S10: Neuroglancer basic welcome and navigation demonstration (associated with Figure S7).**

### Supplementary Text

#### Abbreviations

AAUT: Arm-to-arm U-Tract
ANC: Axial Nerve Cord
APS: Advanced Photon Source
CBC: Cerebrobrachial connective
CBT: cerebrobrachial tract
CNS: Central nervous system
FOV: Field-of-view
FWHM: Full-width half maximum
IFL: Inferior frontal lobe
iLT: Intermediate Longitudinal Tract
oLT: Oral Longitudinal Tract
INC, oINC, aINC: Intramuscular nerve cord (o = oral, a = aboral)
NA: Numerical aperture
OC: Oblique connective
TB: Terabyte

